# High temperature increases centromere-mediated genome elimination frequency in Arabidopsis deficient in cenH3 or its assembly factor KNL2

**DOI:** 10.1101/2022.03.24.485459

**Authors:** Ulkar Ahmadli, Manikandan Kalidass, Lucie Crhak Khaitova, Joerg Fuchs, Maria Cuacos, Dmitri Demidov, Sheng Zuo, Jana Pecinkova, Martin Mascher, Mathieu Ingouff, Stefan Heckmann, Andreas Houben, Karel Riha, Inna Lermontova

## Abstract

Double haploid production is the most effective way of creating true-breeding lines in a single generation. In *Arabidopsis*, haploid induction via mutation of the centromere-specific histone H3 (cenH3) has been shown when outcrossed to wild-type. Here we report that a mutant of the cenH3 assembly factor KNL2 can be used as a haploid inducer. We elucidated that short temperature stress of the *knl2* mutant increased the efficiency of haploid induction from 1 to 10%. Moreover, we have demonstrated that a point mutation in the CENPC-k motif of KNL2 is sufficient to generate haploid inducing lines, suggesting that haploid inducing lines in crops can be identified in a naturally occurring or chemically induced mutant population, avoiding the GMO approach at any stage. In addition, we have shown that the *cenh3-4* mutant, which does not induce haploids under standard growth conditions, functions as a haploid inducer after exposure to short temperature stress.

## Introduction

The haploid generation technology, followed by whole genome duplication, is an effective strategy for accelerating plant breeding, as it allows to obtain true-breeding lines with complete homozygosity in a single step. In the conventional breeding approach, these lines are obtained by inbreeding, and often 7 to 9 generations of inbreeding are performed over several years to achieve the desired level of homozygosity (Britt and Kuppu, 2016). To produce (double) haploids (DHs), two main approaches have been widely used such as the *in vitro* explantation of gametophytic tissues (mainly cultivation of anthers or microspores) and the selective loss of one parental chromosome set *in vivo* through interspecific or intraspecific hybridization (Kalinowska *et al*., 2019). However, depending on the tissue culture or crossability of the species of interest both approaches can only be applied to a limited number of genotypes. Hence, alternative resource-efficient and reliable approaches to produce DHs are strongly required. One way to improve the DH effectiveness is to develop efficient inducer lines that guarantee a high haploid induction rate (HIR) combined with a high-throughput haploid selection system. One promising approach to induce haploids is through centromere-mediated genome elimination (Ravi and Chan, 2010, Kuppu *et al*., 2015).

Centromeres are unique chromosomal regions that mediate the kinetochore protein complex formation and microtubule attachment during cell division (Verdaasdonk and Bloom, 2011, Schalch and Steiner, 2017). Most centromeres are epigenetically defined by nucleosomes containing the centromere-specific histone H3 variant, cenH3 (Allshire, 1997). The cenH3 protein contains two domains, the N-terminal tail, which is a target for post-translational modification, and the C-terminal histone fold domain, which interacts with DNA and other histones to form the nucleosome. The loading of cenH3 to centromeres initiates the assembly of the functional kinetochore complex. The cenH3 loading pathway can be divided into three steps: initiation (centromere licencing), deposition, and maintenance. The centromere licencing factor KNL2, identified in *A. thaliana* showed colocalization with cenH3 throughout the cell cycle except from metaphase to mid-anaphase (Lermontova *et al*., 2013). Furthermore, Sandmann *et al*., (2017) identified a cenH3 nucleosome binding CENPC-k motif of KNL2 at its C-terminal part. The complete deletion of this motif or mutating of its conserved amino acids abolished the localization of KNL2 at centromeres. Thus, it is evident that the CENPC-K motif is functionally required for centromeric localization of KNL2 in *A. thaliana* (Sandmann *et al*., 2017).

Due to its essential function in chromosome segregation, inactivation of cenH3 has been shown to result in chromosome segregation errors and lethality (Ravi and Chan, 2010, Ravi *et al*., 2011). RNAi-mediated knockdown of *cenH3* showed a reduction in its mRNA level (27-43%) and also resulted in a dwarf plant phenotype and meiotic defects in *Arabidopsis* (Lermontova *et al*., 2011). Recently, a mutation in cenH3 named *cenh3-4* has been discovered from the genetic suppressor screen, which increased fertility and promoted meiotic exit in *smg7-6* plants (Capitao *et al*., 2021). The *cenh3-4* is a point mutation (G →A) in the splicing donor site of the 3rd exon of cenH3 showed a reduced amount of cenH3 at centromeres and thus, forming small centromeres. Similar to the cenH3 RNAi transformants, a T-DNA insertion knockout mutant of KNL2 showed a reduced amount of cenH3 at centromeres, decreased growth rate, fertility, and meiotic defects (Lermontova *et al*., 2013), supporting further the functional relationship of both proteins.

Ravi and Chan, (2010) discovered that haploid plants can be obtained by pollination of a *cenh3-1* mutant of *A. thaliana* complemented with a GFP-tail swap construct (fusion of N-terminus of conventional H3 to the C-terminus of cenH3) with different wild-type accessions. This process at the end has resulted in haploid progenies with the genome of the wild-type parent at frequencies as high as 25-45%. If a wild-type female was crossed to a GFP-tail swap male, the proportion of haploid plants was lower. In recent studies, haploids also were achieved by introducing point mutations or small deletions in *Arabidopsis* cenH3 (Karimi-Ashtiyani *et al*., 2015, Kuppu *et al*., 2015, Kuppu *et al*., 2020). Marimuthu *et al*. (2021) showed that cenH3 variants complementing the *cenh3-1* mutant are selectively removed from centromeres during reproduction. Additionally, the authors have demonstrated that the null mutant of VIM1 (VARIANT IN METHYLATION 1) enhances haploid induction frequencies of the complemented *cenh3-1* mutant. The cenH3-based haploid induction approach was successfully extended from *Arabidopsis* to crop plants, but using of homozygous *cenh3* mutant complemented with an altered variant of cenH3 resulted in an average haploid induction frequency below 1% in maize (Kelliher *et al*., 2016). However, recently it has been reported that the use of heteroallelic cenH3 mutation combinations, which are characterized by reduced transmission in female gametophytes, has increased the HIR in maize to 5% (Wang *et al*., 2021). Application of a similar haploid induction approach to wheat resulted in HIR up to 8% (Lv *et al*., 2020).

In accordance with previous studies, an altered cenH3 protein would be sufficient for haploid induction in *Arabidopsis*, but whether an alteration of cenH3 assembly factors such as KNL2 could also be used as a haploid inducer has not been studied yet. Therefore, in this study, we show that a T-DNA knockout mutant of KNL2 is an effective haploid inducer when crossed with *Arabidopsis* wild-type plants. We demonstrate that short-term exposure of *knl2* to heat stress leads to an increase of the haploid induction efficiency from 1% to 10%. Moreover, the stress treatment regime defined for the haploid induction process with the *knl2* mutant also appeared to be effective for the *cenh3-4* mutant. Additionally, we showed that the introduction of a point mutation in the CENPC-k motif of KNL2 is sufficient to create a haploid inducer line.

## Results

### A short temperature stress of the *knl2* mutant increases the efficiency of haploid induction

The T-DNA knockout mutation of the cenH3 loading factor KNL2 (*knl2* mutant) results in a decreased amount of cenH3 protein, suggesting an essential role of KNL2 in the loading of cenH3 at centromeres (Lermontova *et al*., 2013). Therefore, we assumed that the crossing of *knl2* mutant with wild-type *Arabidopsis* might generate haploids similarly to the cenH3 based haploid induction process. To test this hypothesis, the *knl2* mutant was crossed reciprocally with *Arabidopsis* wild-type accession *Landsberg erecta* grown under standard conditions. Flow cytometric analysis (FC) of pools of up to six seeds revealed 1% haploid progeny when *knl2* was used as the female parent (Figure 1, Table 1).

**Figure 1.**
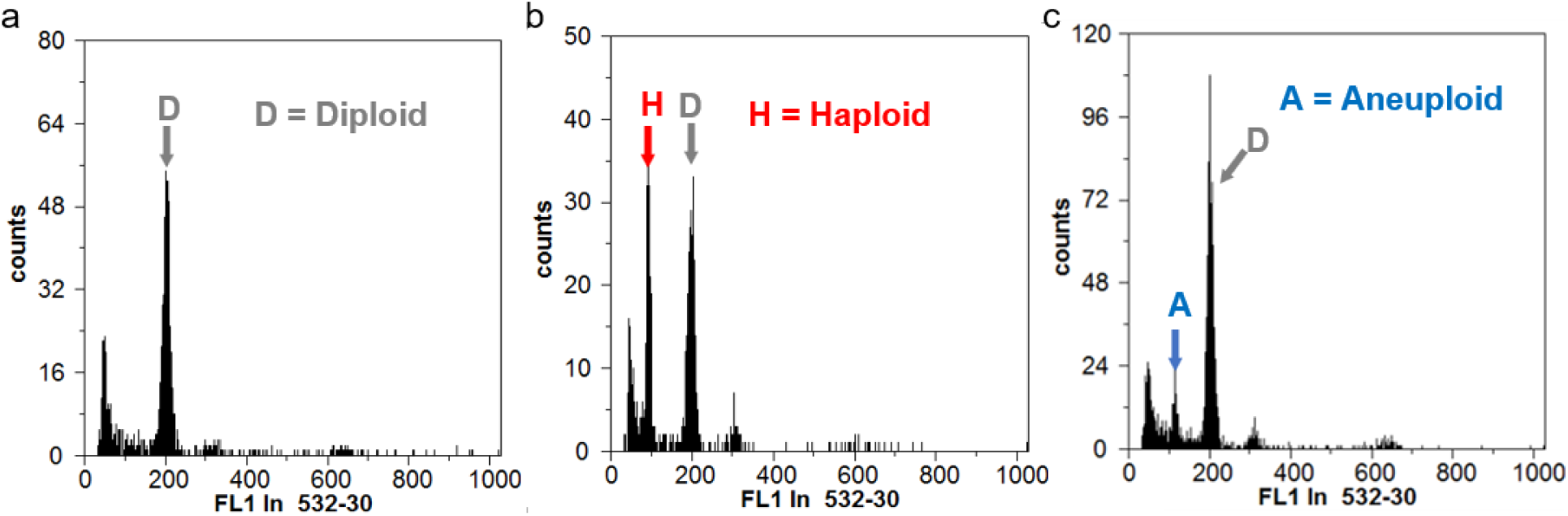
Analysis of haploid seed pools from flow cytometry histograms. a-c Flow cytometry histogram of 6-seed pools containing only diploid (a), haploid and diploid (b), aneuploid, and diploid (c) seeds. The presence of haploid/aneuploid seeds within the pools was determined by evaluating the PI fluorescence intensity on a linear scale. From the seed pools, we considered only one haploid seed per pool compared to the diploid population since the precise number of haploids/aneuploids per pool cannot be measured. Thus, the number of haploids/aneuploids is an underestimation rather than an overestimation.

**Table 1.** Ploidy analysis of seeds derived from the reciprocal crosses of *knl2* with wild-type *A. thaliana* or *gl1-1*.

| Cross (♀ x ♂) | Total seeds | No of haploids | Haploids in (%) | No of aneuploids | Aneuploids in (%) | Conditions |
| --- | --- | --- | --- | --- | --- | --- |
| *knl2* x *Ler* | 196 | 2 | 1 | 1 | 0.5 | Standard condition |
| *Ler* x *knl2* | 335 | 0 | 0 | 0 | 0 | Standard condition |
| *knl2* x *gll-1* | 200 | 0 | 0 | 1 | 0.5 | 25°C |
| ***knl2* x *gll-1*** | **108** | **1** | **0.9** | **1** | **0.9** | **21°C, 400 $\mu\text{mol m}^{-2} \text{sec}^{-1}$** |
| *gll-1* x *knl2* | 126 | 0 | 0 | 0 | 0 | 21°C, 400 $\mu\text{mol m}^{-2} \text{sec}^{-1}$ |
| ***knl2* x *gll-1*** | **256** | **19** | **7.4** | **8** | **5** | **30°C<sup>1</sup>** |
| *gll-1* x *knl2* | 108 | 0 | 0 | 0 | 0 | 30°C<sup>1</sup> |
| *Col* x *gll-1* | 96 | 0 | 0 | 0 | 0 | 30°C<sup>1</sup> |

<sup>1</sup> The plants treated under temperature stress for short time before crossing

Our previous RNAseq data analysis revealed a large number of stress-responsive genes that are differentially expressed in *knl2* seedlings and flower buds compared to wild-type (Boudichevskaia *et al*., 2019). We therefore hypothesized that *knl2* mutant plants may be more sensitive to stress treatment than control plants and that exposure of *knl2* to the stress may increase HIR in its crosses with the wild-type. To support this assumption, the expression of cenH3 and cenH3 assembly factors KNL2, CENP-C, NASP under different stress conditions were analyzed. The gene expression data were retrieved from the *Arabidopsis* transcriptome data platform (http://ipf.sustech.edu.cn/pub/athrdb/). The results showed that these genes were significantly down-regulated in response to various stress treatments like heat, NPA (1-Naphthylphthalamic acid), and fluctuating light (Supplemental Table S1, Figure S1). Thus, increased temperature and light intensity has a strong effect on the expression of cenH3, KNL2 and other key kinetochore components.

To test the impact of stress on *knl2* growth and development and the induction of haploids, it was exposed to either high temperature or high light intensity before crossing. The usage of the *gl1-1* mutant as a crossing partner allows identifying haploids or double haploids based on its trichome-less phenotype (Kuppu *et al*., 2015). First, all plants were cultivated for three weeks under long-day standard conditions (ST) at a temperature of 21/18°C day/night and light intensity at 100 µmol m^-2^sec^-1^. Then, one part of the plants remained under standard growth conditions while others were transferred either to a higher temperature (25/21°C day/night) or high light intensity (400 µmol m^-2^sec^-1^) (Figure 2a). For each growth condition, about 25 *knl2*, 15-20 wild-type, and 15-20 *gl1-1* plants were cultivated. At higher temperatures or light intensity, the phenotypic difference between the wild-type and the *knl2* mutant became more pronounced than under standard growth conditions (Figure S2). Reciprocal crosses were performed between *knl2* mutant plants and *gl1-1* cultivated under growth conditions as described above. The FC analysis of seed pools or trichome less *gl1-1* phenotype analysis of F1 plants revealed no increase in haploid induction efficiency in either type of continuous stress conditions (Table 1 and 2).

**Figure 2.**
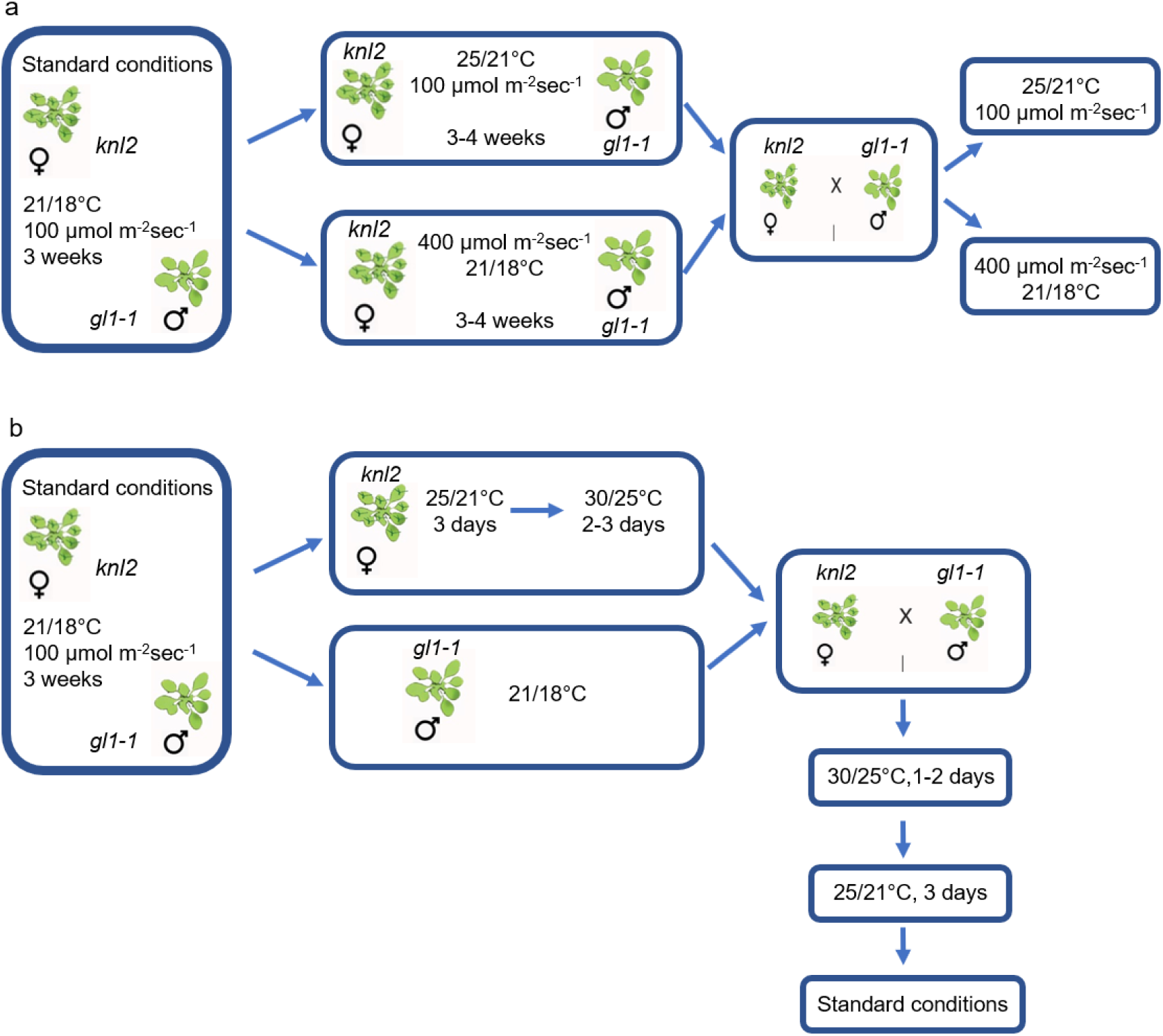
Schematic representation of the crossing of the *knl2* mutant with the *gl1-1* marker line under temperature stress. (a) The *knl2* and *gl1-1* plants were grown under standard growth conditions (21/18°C day/night and 100 µmol m^-2^sec^-1^ light intensity) for three weeks, then the *knl2* mutant plants and *gl1-1* plants were transferred to a growth chamber with constantly increased temperature (25/21°C day/night) or light intensity (400 µmol m^-2^sec^-1^) for 3-4 weeks. The independent crossing of *knl2 and gl1-1* was carried out for increased temperature and light intensities and the plants have remained at the same conditions. (b) Similarly, *knl2* and *gl1-1* plants were grown under standard growth conditions until flowering. Afterward, *knl2* plants were moved to high temperatures (25-21°C day/night) for 3 days followed by short temperature stress (30/25°C day/night) for 2-3 days, while the *gl1-1 L. erecta* marker line was left under standard growth conditions. Then, the *knl2* and *gl1-1* plants were crossed and placed back to 30/25°C day/night for 1-2 days. The temperature was reduced stepwise: first to 25/21°C day/night for three days and then to the standard conditions.

**Table 2.** Phenotype-based selection of plants derived from the reciprocal cross of *knl2* with *gl1-1*.

| Cross (♀ x ♂) | Total no. of plants | No of haploids | Haploids in (%) | Conditions |
| --- | --- | --- | --- | --- |
| *knl2* x *gl1-1* | 144 | 0 | 0 | Standard condition |
| *gl1-1* x *knl2* | 120 | 0 | 0 | Standard condition |
| *knl2* x *gl1-1* | 144 | 0 | 0 | 25°C |
| *gl1-1* x *knl2* | 92 | 0 | 0 | 25°C |
| *knl2* x *gl1-1* | 144 | 1 | 0.7 | 21°C, 400 $\mu\text{mol m}^{-2} \text{sec}^{-1}$ |
| *gl1-1* x *knl2* | 68 | 0 | 0 | 21°C, 400 $\mu\text{mol m}^{-2} \text{sec}^{-1}$ |
| *knl2* x *gl1-1* | 114 | 12 | 10.5 | 30°C<sup>1</sup> |
| *gl1-1* x *knl2* | 29 | 0 | 0 | 30°C<sup>1</sup> |

<sup>1</sup> The plants treated under temperature stress for short time before crossing

Assuming that we did not get an increase in HIR due to adaptation of the *knl2* mutant to continuous stress, the experimental setting was changed and *knl2* mutant plants were exposed to high temperature (30°C) for a short period (2-3 days) before crossing (Figure 2b), while the *gl1-1* crossing partner remained under standard conditions. The temperature was increased and decreased stepwise, as shown in Figure 2. The reciprocal crosses were repeated at least three times in two different growth chambers. FC analysis of seed pools and *gl1-1* mutant phenotype analysis of F1 plants (Figure 3), displayed a similar haploid induction efficiency of 7.4% and 10%, respectively, when heat-stressed *knl2* was used as the female. As the FC analysis of seeds was performed in pools of six seeds and counted only one haploid seed per pool, the number of haploids is compared to the diploid population, which resulted in an underestimation of the number of haploids. No haploids were detected when heat-stressed *knl2* was used as a pollen donor (Table 1 and 2).

**Figure 3.**
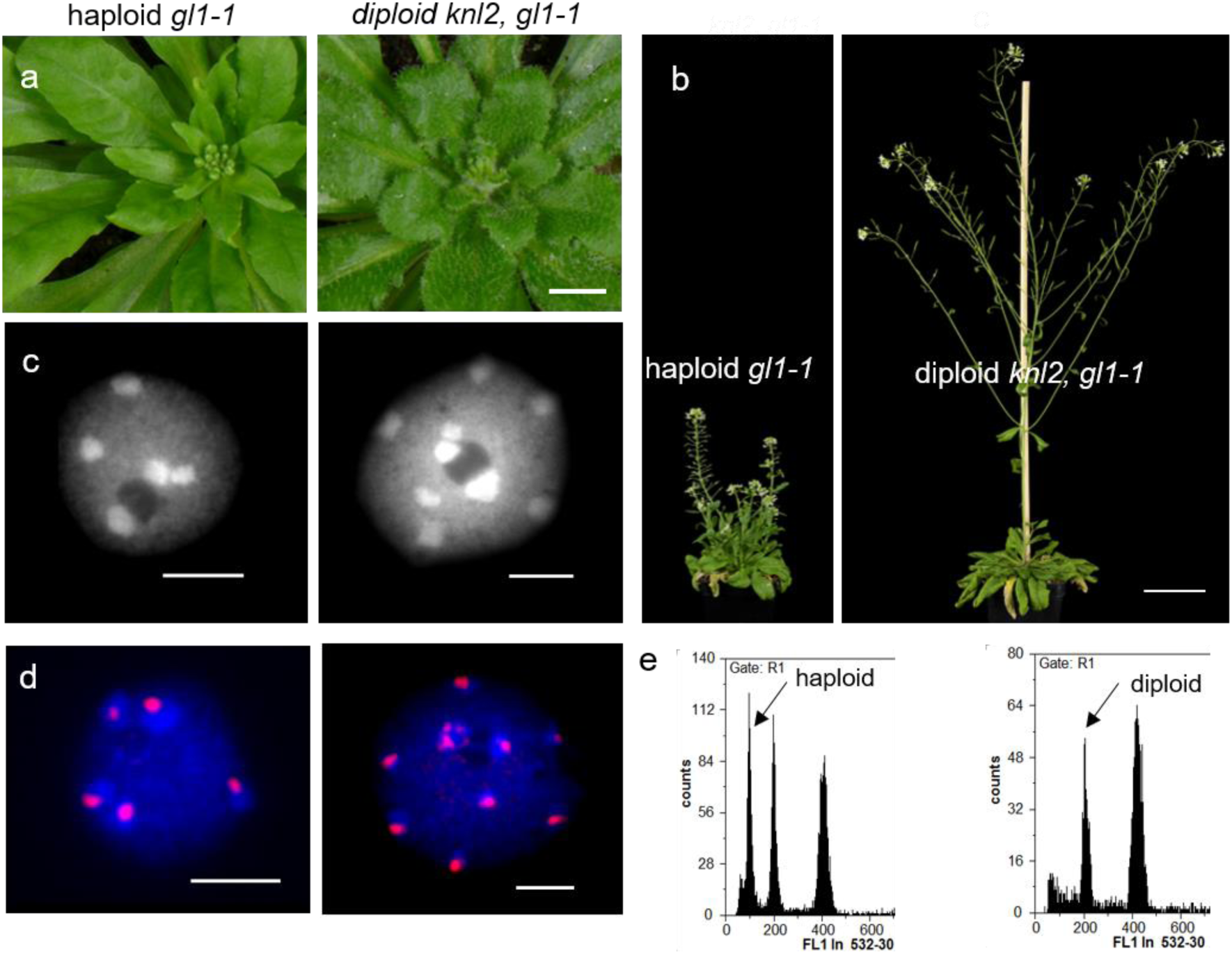
Haploid progenies obtained by genome elimination in crosses of *knl2* with the trichome-less *gl1-1* marker. (a) Comparison of the haploid plant without trichomes (left) and diploid hybrid phenotype with trichomes (right). Scale bar = 1 cm. (b) Phenotype of haploid *gl1-1* and diploid *knl2, gl1-1* hybrid plants during the generative development stage. Scale bar = 5 cm. (c) DAPI stained nuclei isolated from haploid and diploid plants showing a maximum of 5 and 10 heterochromatic chromocenters, respectively. (d) Anti-cenH3 labelled nuclei isolated from haploid and diploid plants showing a maximum of 5 and 10 immunosignals (in red), respectively. The scale bars represent 5 μm. (e) Histogram analysis of nuclei by flow cytometry for a *gl1-1* haploid offspring and control diploid.

The trichomeless plants were much smaller than the corresponding diploids (Figure 3a,b). The 1C nuclei of selected *gl1-1* plants revealed a maximum of 5 chromocenters (Figure 3c) and further, immunostaining of cenH3 also showed 5 chromocenter-localized signals, thus confirming haploidy (Figure 3d). Moreover, a sample flow histogram plot of haploids produced from *knl2* and *gl1-1* crosses and diploid control was shown (Figure 3e).

Besides haploids, 5% of aneuploid seeds were detected by FC (Table 1). However, the control crosses of wild-type Col-0 with *gl1-1 Ler* under the same conditions did not produce haploid plants. Thus, short temperature stress increases the haploidization frequency if *knl2* was used as a female crossing partner.

### A point mutation at the CENPC-k motif of KNL2 results in haploid induction on outcrossing

We previously identified a conserved CENPC-k motif in the KNL2 protein and showed that deletion of this motif or mutagenesis of its conserved amino acids Arg-546 and Trp-555 abolishes the centromeric localization of KNL2 (Sandmann *et al*., 2017). To test whether the introduction of a point mutation into the CENPC-k motif would be sufficient for haploid induction, the genomic KNL2 fragment with the endogenous promoter was cloned into the pDONR221 vector. To substitute the conserved Thr-555 by Arg, PCR-based site-directed mutagenesis was performed (Figure 4). The resulting clone and wild-type KNL2 were subcloned into pGWB640 vector in fusion with EYFP and used for the transformation of the *knl2* mutant. The selected transgenic plants were analyzed for the subcellular localization of KNL2-EYFP fusion protein. In *knl2* mutant complemented with the unmodified KNL2-EYFP construct, fluorescence signals were detected in the nucleoplasm and at chromocenters (Figure 4a), while in *knl2* expressing KNL2-EYFP with the point mutation within the CENPC-k motif, the EYFP signals were detected only in the nucleoplasm (Figure 4b). Immunostaining of root tip nuclei of both variants of transformants with anti-KNL2 antibodies confirmed the centromeric localization of unmutated KNL2 and the nucleoplasmic localization of the variant with a point mutation (Figure 4). Three transgenic lines per construct were selected for the haploid induction experiment under the 30°C degree short temperature stress condition as female crossing partners.

**Figure 4.**
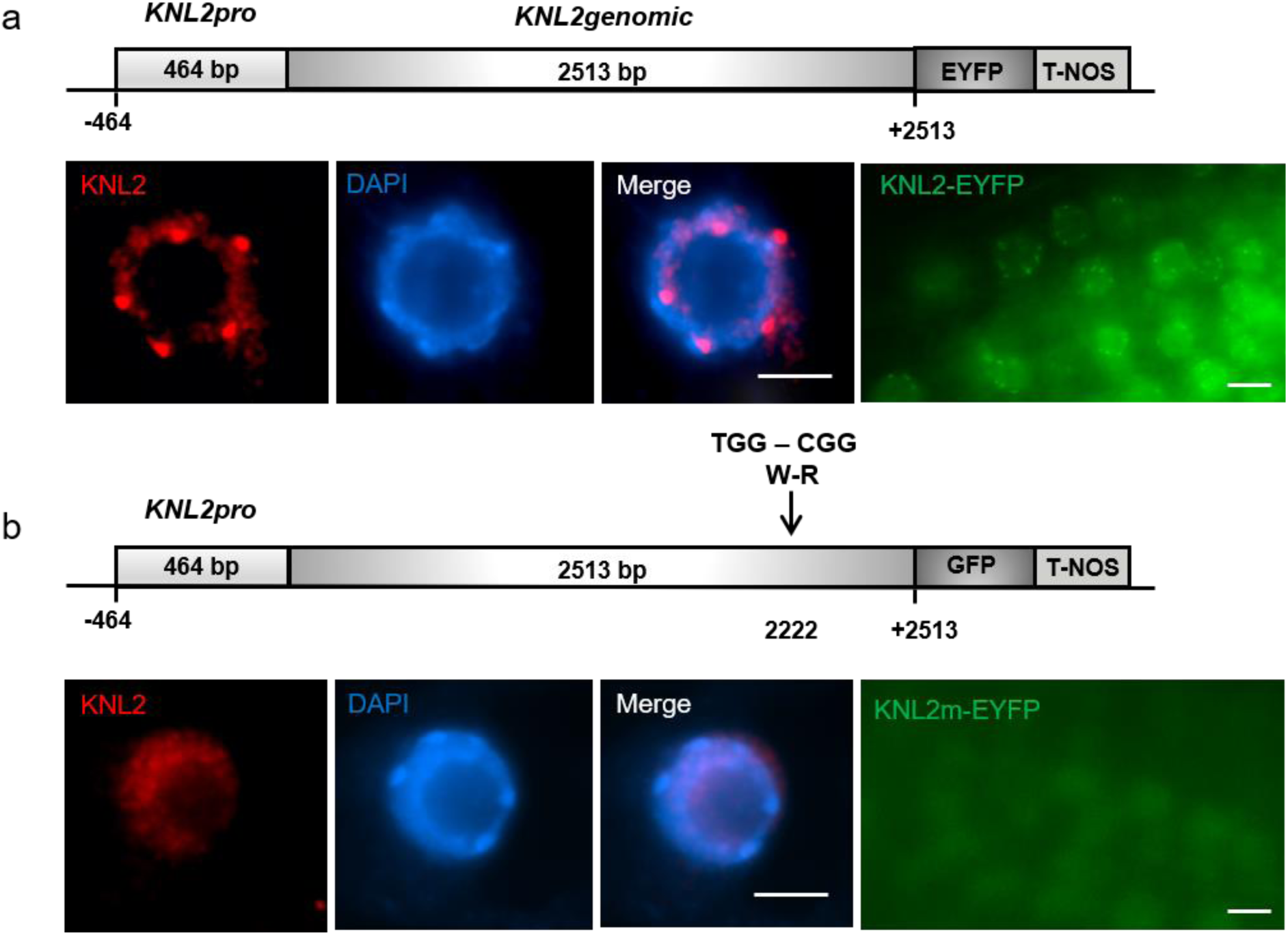
Substitution of the amino acid Trp by Arg within the conserved CENPC-k motif abolished centromeric localization of KNL2. **a-b** Schematic representation of the genomic KNL2-EYFP fusion construct (upper parts) unmodified (a) or carrying the W to R mutation within the CENPC-k motif (b) and the subcellular localization of the corresponding fusion proteins in root tip nuclei of *Arabidopsis* immunostained with anti-KNL2 antibodies (lower parts, panels 1-3) or analyzed by confocal microscopy (lower parts, panel 4). The unmutated KNL2-EYFP fusion protein showed centromeric and nucleoplasmic localization (a) while the variant with point mutation can be detected only in the nucleoplasm (b). The scale bars are 5 µm.

The haploid induction efficiency of the *knl2* mutant complemented by KNL2-EYFP with the point mutation varied from 0.8 to 5.6%. In contrast, no haploids were detected in the case of *knl2* expressing the wild-type KNL2 control construct was used (Table 3).

**Table 3.** Phenotype-based selection of plants derived from the reciprocal cross of *knl2 mutant* with *gl1-1*.

| Cross (♀ x ♂) | Total seeds | No of haploids | Haploids in (%) | No of aneuploids | Aneuploids in (%) | Conditions |
| --- | --- | --- | --- | --- | --- | --- |
| *KNL2gen(W-R) Line 1 x gl1-1* | 126 | 1 | 0.8 | 2 | 1.5 | 30°C<sup>1</sup> |
| *KNL2gen(W-R) Line 2 x gl1-1* | 108 | 6 | 5.6 | 4 | 3.7 | 30°C<sup>1</sup> |
| *KNL2gen(W-R) Line 4 x gl1-1* | 120 | 1 | 0.8 | 2 | 1.7 | 30°C<sup>1</sup> |
| *KNL2genLine1 x gl1-1* | 60 | 0 | 0 | 0 | 0 | 30°C<sup>1</sup> |
| *KNL2genLine2 x gl1-1* | 108 | 0 | 0 | 0 | 0 | 30°C<sup>1</sup> |
| *KNL2genLine2 x gl1-1* | 54 | 0 | 0 | 0 | 0 | 30°C<sup>1</sup> |

<sup>1</sup>The plants treated under temperature stress for short time before crossing

### Analysis of the paternal haploid plants did not reveal any traces of the maternal genome

Next, a PCR-based marker analysis of three double haploid plants was performed to confirm whether only the chromosomes of the pollen donor remained. One genotype-specific marker per chromosome was employed (Figure S3), and in all cases, PCR amplicons were found corresponding to the *gl1-1* mutant. To exclude the presence of small chromosome fragments as a byproduct of the haploidization process as reported by (Tan *et al*., 2015), a Single Nucleotide Polymorphism (SNP) analysis was performed based on Next-Generation Sequence (NGS) reads of DNA samples isolated from three double haploids, one hybrid, and two parental plants. The SNP analysis clearly showed that *gl1-1* double haploid plants do not contain any residues of the maternal *knl2* chromosome complement (Figure 5). The hybrid, by contrast, was heterozygous throughout its genome. A read depth analysis did not show any chromosomal aberrations as observed by (Tan *et al*., 2015)(Figure S4).

**Figure 5.**
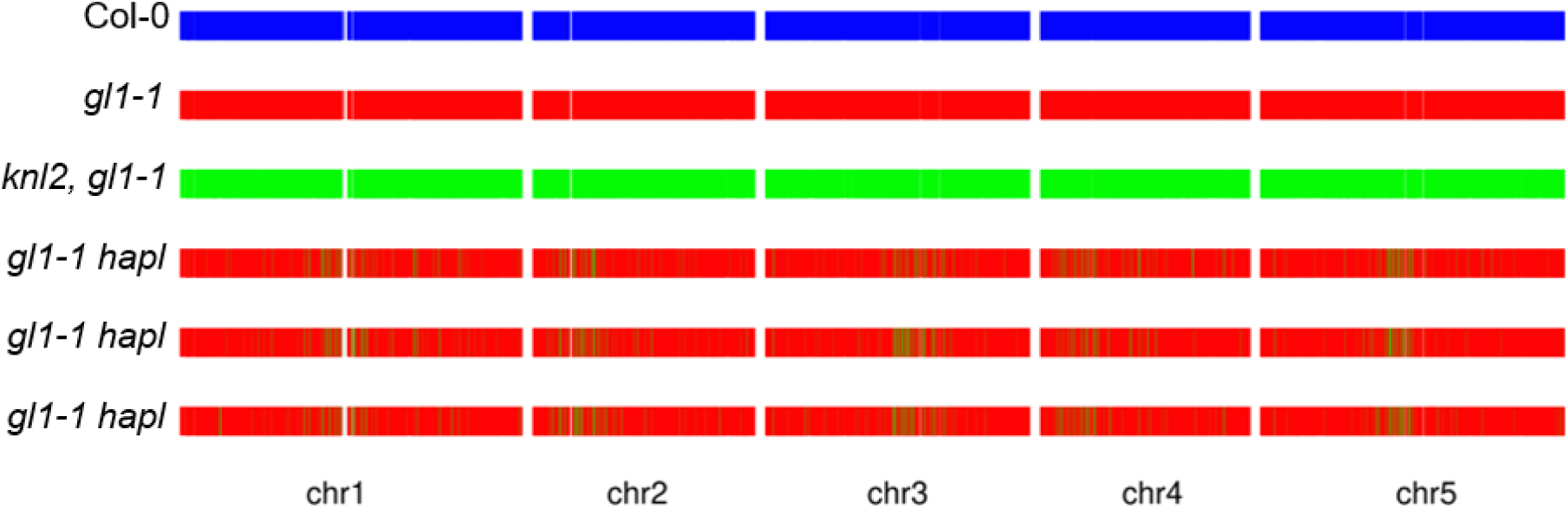
Confirmation of haploid progeny obtained by crossing of *knl2* with the trichome-less *gl1-1* mutant. Single Nucleotide Polymorphism (SNP) analysis of three *gl1-1* double haploids, *knl2, gl1-1* hybrid, *gl1-1*, and Col plants. The results displayed that the hybrid plants were completely heterozygous whereas *gl1-1* double haploid plants do not contain any residues of the maternal *knl2* mutant genome.

### Plants exposed to high-temperature show reduced seed setting as a result of increased mitotic and meiotic abnormalities

In *Arabidopsis* wild-type and *knl2* mutant, exposure to high temperature (30°C) using the regime indicated (Figure 2b) resulted in decreased seed setting and an increased number of aborted seeds after selfing. However, this effect was more pronounced in *knl2* mutant compared to wild-type (Figure 6). Thus, after exposure to high temperature, the average seed number per silique was reduced from 53 to 33 in Col and 45 to 14 in *knl2* mutant, while the number of aborted seeds was increased from 2 to 4 in Col and 5 to 9 in *knl2* mutant, respectively. To address whether the reduction in fertility was based on defects during meiosis, male meiotic chromosome spread analysis was performed in wild-type and *knl2* plants exposed to the same growth conditions as mentioned above.

**Figure 6.**
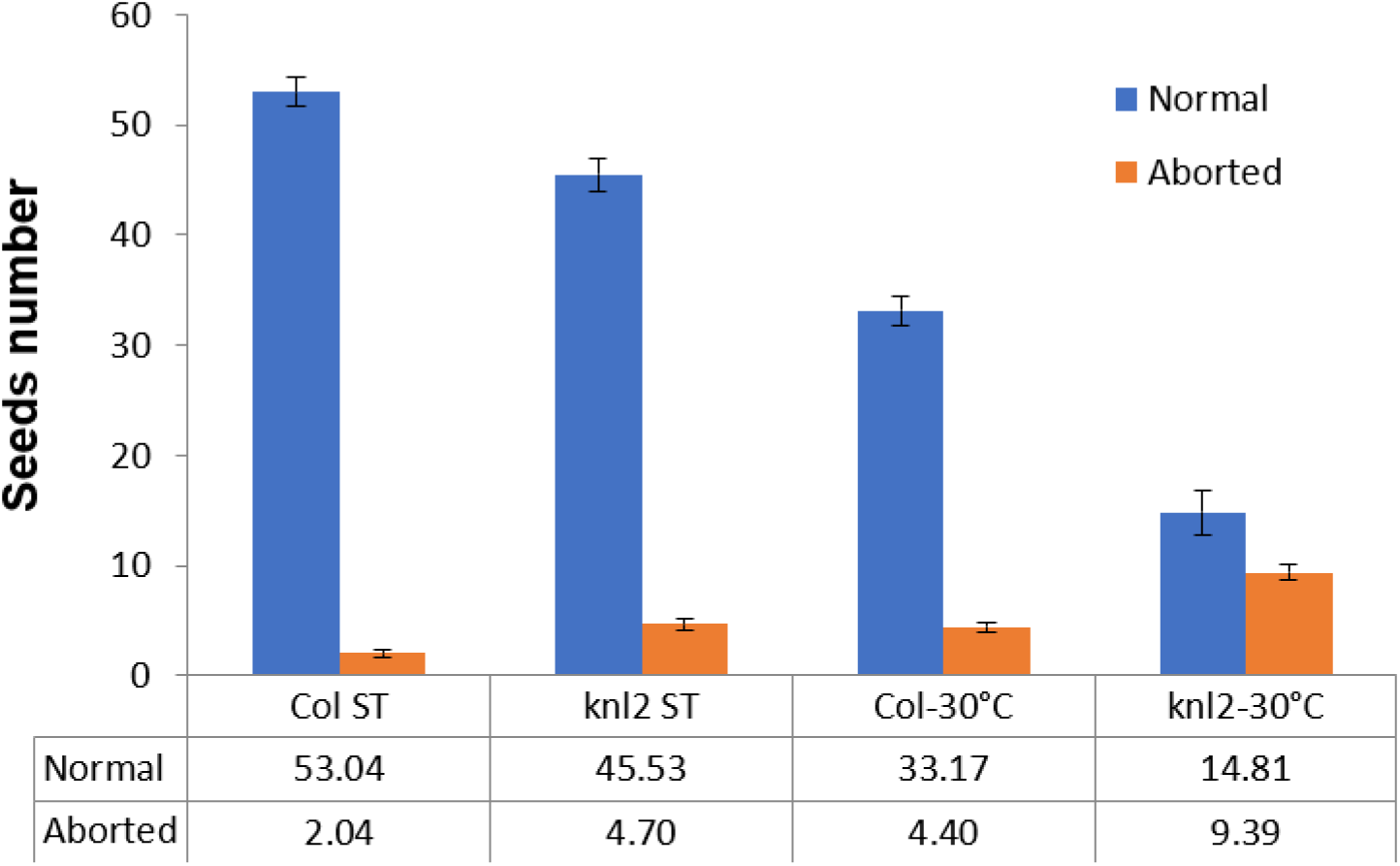
Exposure of *Arabidopsis knl2* mutant and wild-type to high temperature resulted in a decreased seed setting and an increased number of aborted seeds. Seed setting analysis was performed on selfed plants either continuously grown under standard growth conditions or exposed for 4 days to 30/25°C (day/night) as it is indicated in Figure 2b.

No meiotic defects were observed neither at 21°C nor at 30°C (two plants each) in wild-type plants, i.e. homologous chromosomes undergo synapsis at pachytene, five bivalents are inevitably found at metaphase I, homologous chromosomes segregate during the first meiotic division, and the sister chromatids are separated during the second meiotic division (Figure 7). In *knl2* plants grown at 21°C (2 out of 4 plants) and 30°C (4 out of 4 plants), meiotic defects were detected including synapsis defects (asynapsis and interlocks), as well as lagging chromosomes and chromosome fragmentation during the first and second meiotic divisions (Figure 7).

**Figure 7.**
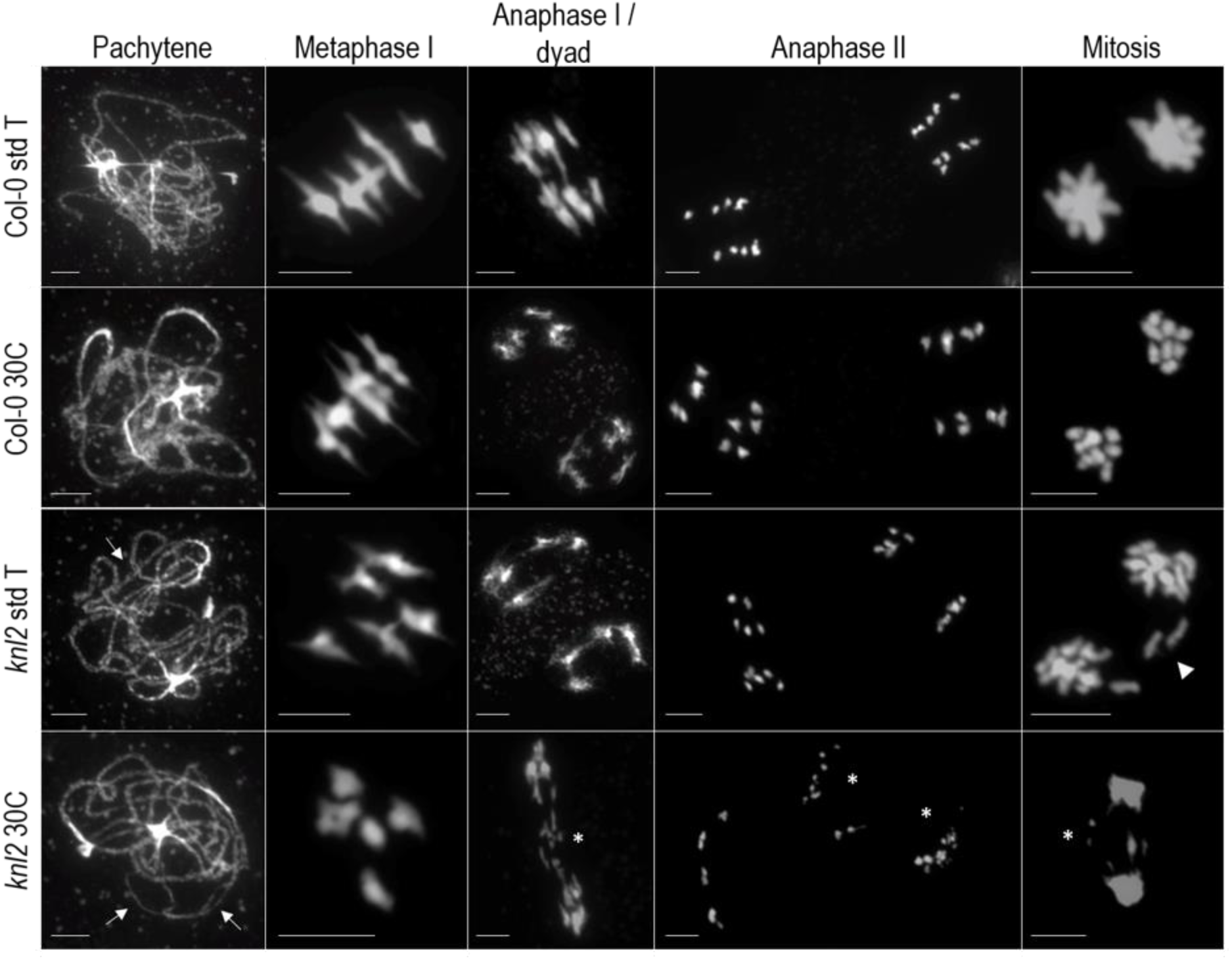
Mitotic and meiotic defects under high temperature in the *knl2* mutant. Male meiosis and mitosis (from tapetum cells) in wild-type and knl-2 plants grown at 21°C or exposed to 30°C as it is indicated in Figure 2b. In wild-type plants grown at either temperature show no errors in meiosis and mitosis progress. During meiosis, homologous chromosomes undergo synapsis at pachytene, form five bivalents at metaphase I, segregate to opposite poles at anaphase I/dyad, and separate sister chromatids during anaphase II. During mitosis, sister chromatids separate to opposite poles. In knl2 plants, meiotic and mitotic defects are found, including asynapsis and interlocks during pachytene (arrows), lagging chromosomes (arrowhead), and chromosome fragmentation (asterisks) during mitotic and meiotic divisions. The chromosomes were stained with DAPI (blue). The bar represents 5 µm.

The degree of observed defects varied among plants and was more pronounced at 30°C than at 21°C (Figure 8). Due to observed meiotic chromosome fragmentation, mitotic divisions of tapetum cells from the same plants were studied. Similar to meiosis, mitotic defects were observed in *knl2* plants grown at 21°C (2 out of 4 plants) and 30°C (3 out of 4 plants), which include lagging chromosomes, anaphase bridges, and chromosome fragmentation (Figure 7) and were more pronounced at a higher temperature. No obvious mitotic defects were found in wild-type grown at either temperature (two plants each). In a nutshell, mitotic and meiotic defects were found to varying degrees in *knl2* plants that were more frequent in plants grown at 30°C, while no noticeable defects were observed in wild-type plants at both temperature regimes.

**Figure 8.**
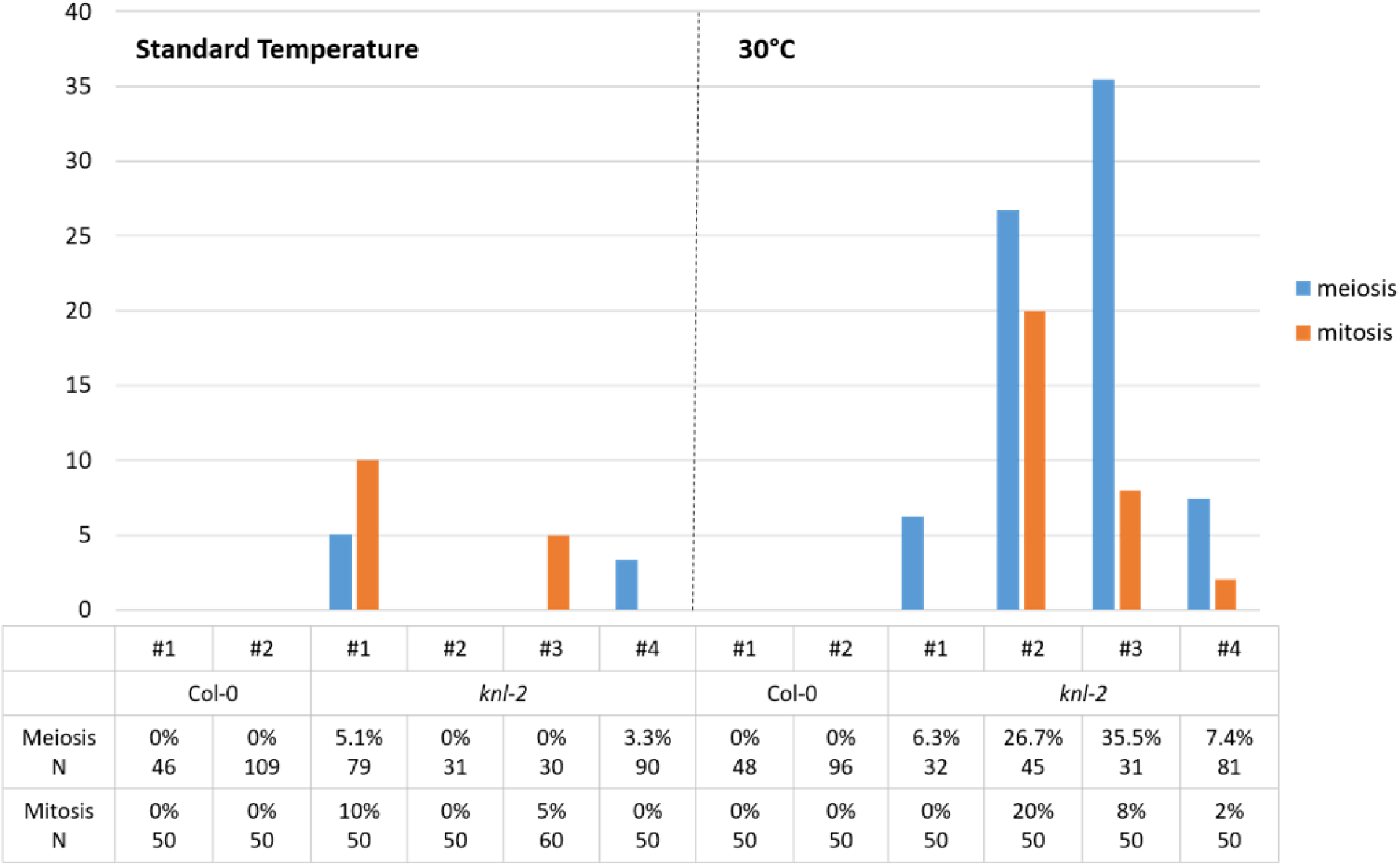
Meiotic and mitotic phenotype of independent wild-type and *knl-2* plants at 21°C and 30°C. The percentage of observed cells with defects per independent plant (meiosis: synapsis defects, lagging chromosomes, and chromosome fragmentation; mitosis: lagging chromosomes, anaphase bridges, and chromosome fragmentation) and the number of cells analyzed were indicated.

### A *cenh3-4* mutation induces haploid formation under temperature stress

The *cenh3-4*, a point mutation of cenH3 (G →A amino acid substitution in the third exon of cenH3) showed a substantially reduced level of cenH3 at the centromere and causes defects in the mitotic spindle. Nevertheless, the reduced level of cenH3 induced haploid plants only with a very low frequency (0.2%) when crossed with wild-type plants. Thus the smaller centromere size was not efficient to trigger haploidization in *Arabidopsis* (Capitao *et al*., 2021). Considering the effect of heat stress on the *knl2* mutant, we tested the haploid induction rate with *cenh3-4* mutants cultivated under heat stress. In our initial experiment, *cenh3-4* plants were exposed to increasing temperatures for a longer period before pollination, but transferred to standard conditions (22°C) immediately after the pollination. The *cenh3-4* mutant and trichome-less *gl1-1* plants were grown under 22°C for two weeks, plants at the earlier rosette stage were transferred to the chambers containing different temperatures varying from 16ºC to 30ºC and cultivated for additional two to three weeks until they formed flowers (Table 4). Then, the *cenh3-4* mutant plants were pollinated with pollen from trichome-less *gl1-1* plants grown under 22°C, and the pollinated plants were transferred to 22°C. The haploid induction rate was determined based on the trichome-less phenotype. Out of 181 progeny plants grown at 26ºC and 253 plants grown at 30ºC, one and two haploid plants were identified, respectively (Table 4). No haploid plants were found among more than 3197 progeny plants at lower temperatures (16, 22, 24°C). This data supports the notion that increased temperature promotes haploid induction in centromere-impaired plants.

**Table 4.** Analysis of haploid induction in cenH3 mutants based on phenotype and ploidy levels by flow cytometry.

| Cross (♀ x ♂) | Total seeds/plants | No of haploids | Haploids in (%) | No of aneuploids | Aneuploids in (%) | Conditions |
| --- | --- | --- | --- | --- | --- | --- |
| *cenh3-4* x *gl1-1* | 1023 | 0 | 0 | - | - | 16°C <sup>1</sup> |
| *cenh3-4* x *gl1-1* | 998 | 0 | 0 | - | - | 22°C <sup>1</sup> |
| *cenh3-4* x *gl1-1* | 1176 | 0 | 0 | - | - | 24°C <sup>1</sup> |
| ***cenh3-4* x *gl1-1*** | **181** | **1** | **0.55** | - | - | **26°C <sup>1</sup>** |
| *cenh3-4* x *gl1-1* | 86 | 0 | 0 | - | - | 28°C <sup>1</sup> |
| ***cenh3-4* x *gl1-1*** | **253** | **2** | **0.79** | - | - | **30°C <sup>1</sup>** |
| ***cenh3-4* x *gl1-1*** | **336** | **11** | **3.3** | **9** | **2.7** | **30°C <sup>2</sup>** |
| *gl1-1* x *cenh3-4* | 90 | 0 | 0 | 1 | 1.1 | 30°C <sup>2</sup> |
| ***cenh3-4* x *gl1-1*** | **96** | **4** | **4.1** | - | - | **30°C <sup>3</sup>** |
| *cenH3 RNAi* x *gl1-1* | 120 | 0 | 0 | 0 | 0 | 30°C <sup>2</sup> |
| *gl1-1* x *cenH3 RNAi* | 66 | 0 | 0 | 0 | 0 | 30°C <sup>2</sup> |

<sup>1</sup>*cenh3-4* plants treated under temperature stress continuously and analyzed based on phenotypic differences
<sup>2</sup>*cenh3-4* plants treated under temperature stress shortly before crossing and analyzed by flow cytometry
<sup>3</sup>*cenh3-4* plants treated under temperature stress shortly before crossing and analyzed based on phenotypic differences

Nevertheless, the HIR in *cenh3-4* plants at continuous heat stress regime was substantially lower than the one achieved with *knl2* mutants using heat treatment described in Figure 2a. Therefore, we recapitulated the experiment using the same conditions as for the *knl2* plants. The *cenh3-4* plants were subjected to 30°C for 2 days (Figure 2b), then pollinated with *gl1-1* pollen, cultivated for additional 2 days at 30°C followed by 3 days at 25°C before transferring to standard conditions. Analysis of 56 seed pools (6 seeds per pool) with a total of 336 seeds by flow cytometry revealed a haploid and aneuploid induction rate of 3.3% and 2.7%, respectively (Table 4). Moreover, the haploid induction frequency of 4.1 % was determined based on the trichomeless phenotype of *gl1-1* (Table 4). This result indicates that sustaining the temperature stress for several days after pollination further improves HIR.

Additionally, cenH3 RNAi transformants that revealed a substantial reduction of cenH3 at centromeres (Lermontova *et al*., 2011) were tested as haploid inducers in combination with heat stress. Reciprocal crosses of RNAi with *gl1-1* were performed under short-term 30°C-stress conditions (Figure 2b). In contrast to *cenh3-4*, FC analysis of seeds did not reveal either haploids or aneuploids (Table 4).

## Discussion

In most eukaryotes, kinetochore assembly is primed by cenH3 and multiple kinetochore protein complexes are required for accurate chromosome segregation. We showed that the disruption of the cenH3 loading machinery via the inactivation of the centromere licensing factor KNL2 of *Arabidopsis* resulted in the generation of haploids (HIR – 1%) on outcrossing with wild-type. To enhance the efficiency of haploid induction, the *knl2* mutant plants were subjected to various stress conditions, such as increased temperature and light intensity as we found that deregulated expression of KNL2 leads to differential expression of many stress-responsive genes (Boudichevskaia *et al*., 2019). However, cultivation of *knl2* plants under long-term stress conditions (25/20°C day/night, 100 µmol m^-2^sec^-1^ light intensity or 21/18°C day/night and 400 µmol m^-2^sec^-1^ light intensity) did not increase HIR in the reciprocal crosses of *knl2* with wild-type. Assuming that the applied cultivation regimes either did not cause severe stress or that the plants had adapted to the long-term treatment, short-term treatment with high temperature (30/25°C day/night) has been applied for 2-3 days prior to the crossing experiments. Indeed, exposure of *knl2* to high temperatures for a short period allowed us to increase the HIR to up to 10 %.

Moreover, the same heat stress treatment applied to the *cenh3-4* mutant (Capitao *et al*., 2021) resulted in an increase in HIR from 0.2% under standard conditions to 4.1%. Interestingly, our data also show that the more efficient HIR is achieved when heat stress is prolonged to the postfertilization period. This indicates that the haploid induction in centromere impaired mutants is conditioned by temperature stress during both ovule development as well as early embryogenesis. Originally, it was thought that haploids could only be obtained by crossing the *cenh3* mutant lines complemented by a modified version of cenH3 with the wild-type, as the elimination of one of the genomes is caused by competition between two structurally different cenH3 variants for the deposition to centromeres (Ravi *et al*., 2014, Thondehaalmath *et al*., 2021). To understand the mechanism of genome elimination in such crosses, Marimuthu *et al*. (2021) analyzed the distribution of the altered cenH3 (GFP-tail swap) variant in gametes and at different developmental stages of hybrid zygotes. It has been shown that altered cenH3 is selectively removed from mature *Arabidopsis* eggs and early hybrid zygotes while at the later zygotic stages, cenH3 and GFP-tail swap preferentially can be loaded into the centromere of the wild-type parent, whereas the cenH3-depleted mutant chromosomes are not able to reconstitute new cenH3 chromatin and undergo elimination. However, Wang *et al*. (2021) and Lv *et al*. (2020) have demonstrated that heterozygous cenH3 mutants of maize and wheat, respectively, can also function as efficient haploid inducers in crosses with the wild-type in both directions, despite the lack of competition between the two structurally different cenH3 variants. In these cases, it was assumed that weak centromeres are formed due to cenH3 dilution that occurs as a result of postmeiotic divisions in gametogenesis. Since female haploid spores undergo three mitotic divisions and male only two, the level of cenH3 in female gametes is expected to be lower (Wang *et al*., 2021). When a heterozygous *cenh3-1* mutant of *Arabidopsis* was used as an HI in crosses with wild-type, haploids could be generated at a frequency of ∼1%, indicating that also in *Arabidopsis* haploids can be produced without alteration of cenH3 (Marimuthu *et al*., 2021).

Using *knl2* and *cenh3-4* mutants of *Arabidopsis*, we further demonstrated that a competition of structurally different variants of cenH3 is not always a prerequisite for haploid induction. Thus, similar to maize and wheat, the centromere size model of (Wang and Dawe, 2018) can be applied to the haploid induction approach based on *knl2* and *cenh3-4* mutants. We suggest that applying temperature stress to *knl2* and *cenh3-4* mutants can weaken their centromeres additionally compared to standard growth conditions and therefore lead to an increase in HIR (Figure 9).

**Figure 9.**
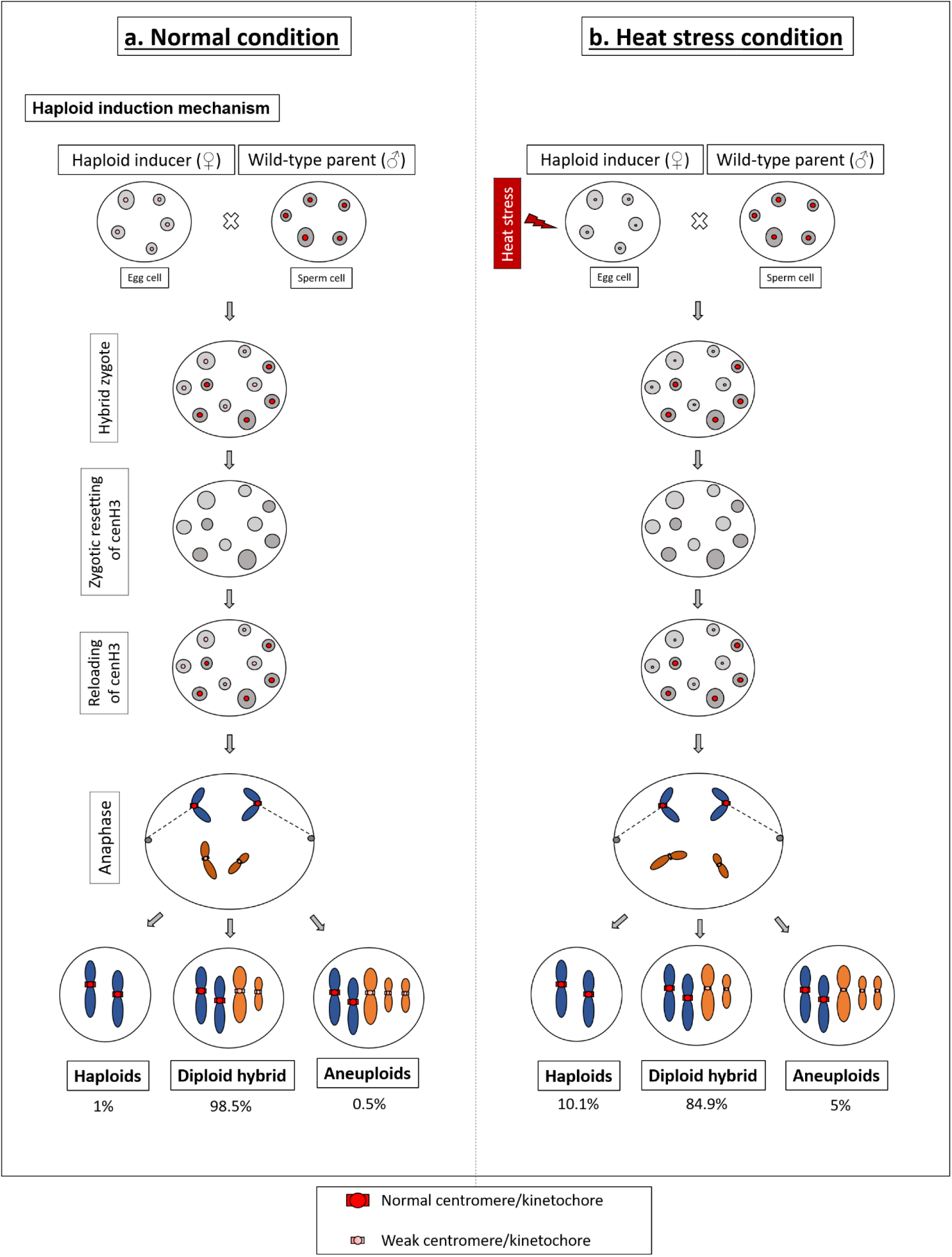
A model comparing a uniparental chromosome elimination mechanism in crosses of the *knl2* mutant × wild-type under standard (a) and heat stress condition (b). Model explains the elimination of uniparental chromosomes in haploid inducer *knl2* mutant crossed with wild-type under standard condition (left panel) and heat stress condition (right panel). (a) In standard conditions, the combination of small size centromeres of the haploid inducer (*knl2* mutant) with ‘normal’ wild-type centromeres in the hybrid zygote leads to centromere competition, followed by complete or partial elimination of the genome and formation of haploid (1%) and aneuploid (0.5%) progeny of wild-type, respectively. (b) Under heat stress conditions, the haploid inducer (*knl2*) chromosomes have severe mitotic and meiotic defects as shown in (Figure 7), which leads to the centromere inactivity and making very small centromeres. Therefore, the haploid induction rate was increased to 10% when heat stressed *knl2* mutant crossed with wild-type plants.

The bioinformatic analysis revealed a reduced expression of genes coding for the kinetochore proteins such as cenH3, KNL2, and CENP-C under stress conditions. And while this reduction is not critical in the wild-type, in *knl2* and *cenh3-4* it amplifies the effect of mutations. This suggestion can be supported by our data showing that the frequency of mitotic and meiotic defects in *knl2* can be increased after short-term heat stress treatment while wild-type plants cultivated under the same conditions did not show any mitotic or meiotic abnormalities. Our RNAseq data analysis has revealed that a high number of transposable elements was activated in seedlings and flower buds of the *knl2* mutant cultivated under standard growth conditions (Boudichevskaia *et al*., 2019). Thus, it can be assumed that exposure of *knl2* to heat stress might result in even an increased number of active transposons compared to the standard conditions and disturb chromatin organization consequently (Probst and Mittelsten Scheid, 2015). These are some of the possibilities behind the temperature stress-induced haploid formation but, a clear mechanism is still needed to be elucidated.

It is important to understand why haploids cannot be obtained when heat-treated mutants are used as pollen donors. There are relatively few examples in which the effects of temperature stress on female reproductive organs have been investigated, but much more is known of the effects of temperature stress on male reproductive structures (Hedhly *et al*., 2009). Using tomato male-sterile and male-fertile lines, (Peet *et al*., 1998) have demonstrated that stress applied to the pollen donor plant before and during pollen release decreased seed number and fruit set more severely than heat stress applied to the developing ovule. Thus, heat stress treatment of *knl2* and *cenh3-4* mutants as pollen donors can lead to a decrease in pollen viability and the inability to fertilize the egg. At the same time, fertilization of the ovule with viable pollen will not lead to the process of genome elimination. In contrast, heat stress treatment of mutants as maternal crossing partners can only lead to a weakening of the centromeres but does not affect the viability of the ovules.

The most notorious phenotypes in the *knl2* mutant plant’s meiosis were the presence of lagging chromosomes and fragmentation. Lagging chromosomes could appear as a consequence of weaker or defective centromere activity in *knl2* plants. Fragmentation appeared during mitosis and meiosis. Interestingly, two plants grown at 30°C showed fragmentation in anthers during the first meiotic division, and the other two plants showed during the second meiotic division. This suggests that fragments could be originated by different mechanisms. One possibility is that interlocks observed during pachytene are not properly repaired and another would be the defective DNA repair in pathways involving both inter-sister and inter-homolog events. Besides, anaphase bridges were also observed due to the effect of heat stress in the *knl2* mutant. In a recent publication describing the comparative analysis of seedlings and flower bud transcriptomes of *knl2* mutant and wild-type, it has been shown that genes coding for proteins involved in DNA repair were overrepresented among down-regulated *knl2* genes (Boudichevskaia *et al*., 2019).

For instance, the down-regulated genes were KU70 (AT1G16970), KU80 (AT1G48050), and LIGASE 4 (AT5G57160), the key players participating in the canonical non-homologous end joining, RAD51 (AT5G20850), essential for meiotic repair of DSBs caused by AtSPO11-1 (Li *et al*., 2004), DMC1 (AT3G22880), known to promote interhomolog recombination, SMC6A (At5G07660) and SMC6B (At5G61460), two components of the SMC5/6 complex, engaged in DNA repair, meiotic synapsis, genome organization, and stability. Previously it was shown that high temperatures disturb genome integrity by causing strand breakages and impending DNA repair (Kantidze *et al*., 2016) and crosstalk between heat stress and genotoxic stress in *Arabidopsis* has been demonstrated Han *et al*. (2020). We speculate that because of reduced expression of genes encoding components of the DNA repair mechanism, the *knl2* mutant cannot cope with heat-induced DNA damage as efficiently as the wild-type and therefore increased mitotic and meiotic defects in *knl2* after exposure to high temperature has been detected (Figure 10).

**Figure 10.**
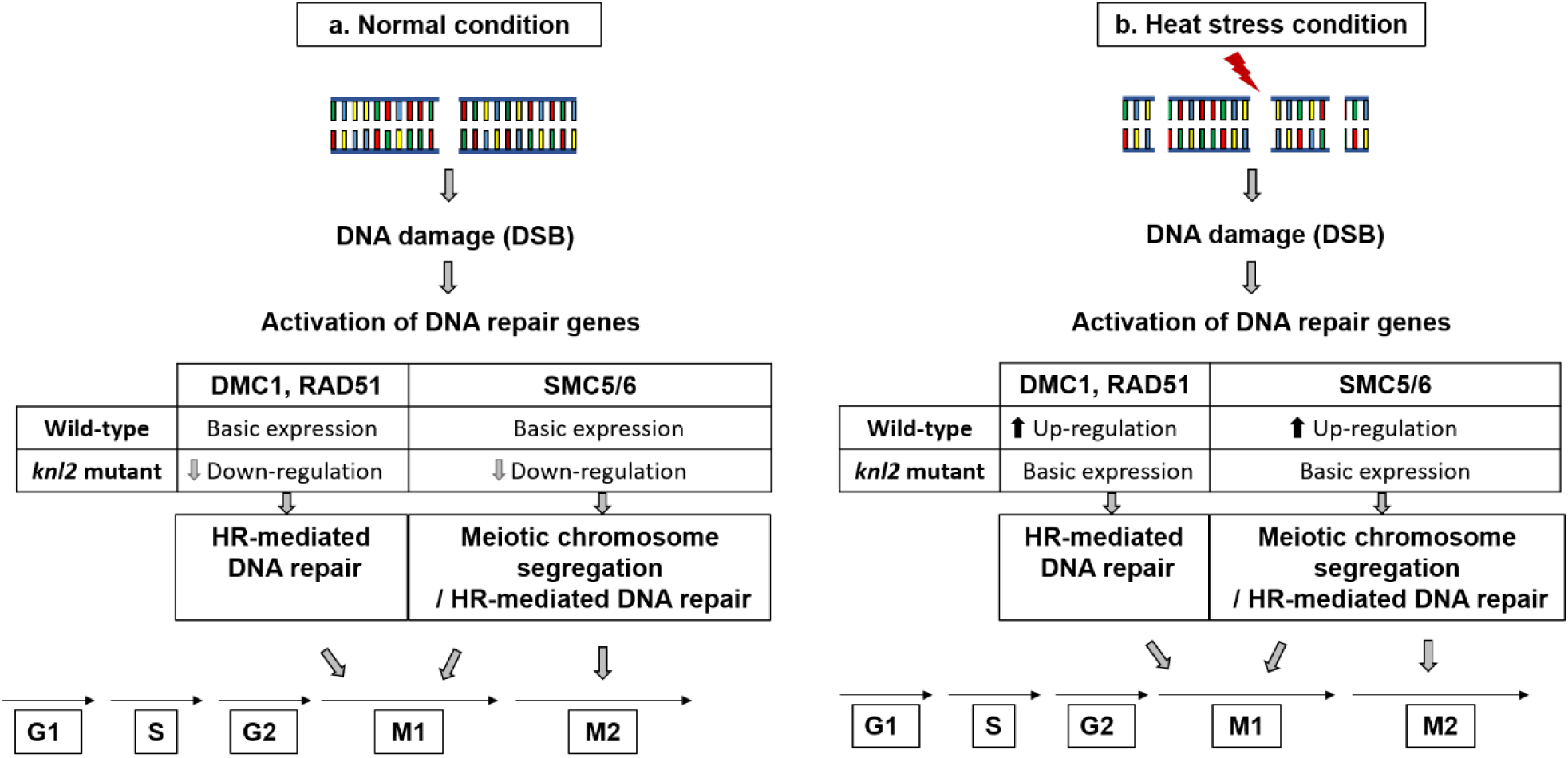
Effect of DNA repair pathway in wild-type and *knl2* mutant under normal (a) and heat stress conditions (b) In standard growth conditions (left panel), *knl2* mutant shows a reduced expression of genes encoding the DNA repair proteins (Boudichevskaia *et al*., 2019) that might correlate with the mitotic and meiotic abnormalities in the mutant. Exposure of *knl2* and wild-type to heat stress (right panel) results in increased DNA damage. In wild-type, this is accompanied by increased expression of DNA repair genes (Han *et al*., 2020), whereas in *knl2* these genes are down-regulated under standard growth conditions and therefore, their expression under heat stress cannot be sufficiently increased. In the *knl2* mutant, heat stress increases DNA damage, mitotic and meiotic abnormalities, and reduced expression of cenH3 and other kinetochore related genes. These factors could be the reason for the increase in HIR when heat stressed *knl2* mutant used as a haploid inducer.

In principle, we can expect that the cenH3 RNAi transformants with strongly reduced cenH3 levels can also work as efficient haploid inducers (Lermontova *et al*., 2011). However, subjecting cenH3 RNAi transformants to heat stress did not result in haploid formation when crossed with the untreated wild-type. Previously, it has been shown that the level of cenH3 in the cenH3 RNAi transformants was strongly reduced in leaves than in root tips enriched in meristematic cells (Lermontova *et al*., 2011). Based on previously published data, we hypothesized that this may be due to a decreased activity of the CaMV 35S promoter (Holtorf *et al*., 1995) and suppression of post-translational gene silencing in meristems, which may cause the ineffective function of the RNAi machinery in these tissues (Mitsuhara *et al*., 2002). Using maize cenH3 RNAi lines complemented by the *AcGREEN-tail swap-CENH3*, Kelliher *et al*. (2016) have demonstrated that in crosses with wild-type these lines can generate 0.24% maternal and 0.07% paternal haploids. Thus, the method of obtaining haploid inducers through reduction of cenH3 levels in plants by expressing cenH3 RNAi constructs appeared to be inefficient.

The introduction of point mutations into cenH3 and the conserved CENPC-k motif of KNL2 has shown to be sufficient to generate haploid inducer lines. Thus, *knl2* and *cenh3* mutants for crop species can be obtained via the chemical ethyl methanesulfonate (EMS) mutagenesis to avoid using transgenic plants at all steps of haploid production. Alternatively, mutants can be produced by targeted mutagenesis using the CRISPR-Cas9 approach. In either case, complementation with altered cenH3 variants is not required, making the production of haploid inducers much easier. Moreover, under standard growth conditions, the growth rate of the *cenh3-4* mutant is similar to that of the wild-type, while the growth rate of *knl2* is slightly reduced. Thus, we believe that obtaining vigorous haploid inducers and short-term exposure of them to heat stress before crossing with the wild-type has great potential for application in plant breeding.

## Acknowledgements

This research was supported by the German Federal Ministry of Education and Research (Plant 2030, Project 031B0192NN, HaploTools) and by the Deutsche Forschungsgemeinschaft (LE2299/3-1; LE2299/5-1) by the European Regional Development Fund-Project ‘REMAP’ (No. CZ.02.1.01/0.0/0.0/15_003/0000479) to KR. We thank Heike Kuhlmann and Oda Weiss for excellent technical assistence. We also acknowledge the Plant Sciences Core Facility of CEITEC MU for support with plant cultivation.

## Materials and Methods

### Plasmid construction, plant transformation, and plant growth conditions

To analyze whether complementation of the *knl2* mutant with the genomic KNL2::KNL2-EYFP fusion construct would abolish the ability of *knl2* to induce haploids, genomic KNL2 fragment (464 up to +2513 relative to the transcriptional KNL2 start site) was amplified by PCR from Col-0 genomic DNA using KNL2-attB1gensh and KNL2-attB2 primers (Supplemental Table S2) and cloned into the pDONR221 vector. These constructs were used to generate KNL2:KNL2-EYFP fusion construct using the pGWB640 vector (https://novoprolabs.com/vector/Vgy4dmna). The substitution of conserved amino acid Trp by Arg within the CENPC-k motif of KNL2 was performed by PCR using the Phusion site-directed mutagenesis kit (Thermo Fisher Scientific). A KNL2::KNL2-EYFP/pDONR221 construct was PCR mutagenized using the following primer pairs: KNL2gen_W_R_f and KNL2gen_W_R_r for the substitution of Trp by Arg (Supplemental Table S2). *Arabidopsis thaliana* plants were transformed according to the flower dip method (Clough and Bent, 1998). T1 transformants were selected on Murashige and Skoog medium containing 20 mg L-1 phosphinothricin (PPT). The plants were propagated under short-or long-day conditions in a cultivation room at 8 h light/20°C:16 h dark/18°C and 16 h light/20°C:8 h dark/18°C, respectively.

### Immunostaining and microscopy analysis of fluorescent signals

Immunostaining of nuclei/chromosomes was performed as described previously (Sandmann *et al*., 2016). Wide-field fluorescence microscopy was used to evaluate and image the nuclei preparations with an Olympus BX61 microscope (Olympus, Tokio, Japan) and an ORCA-ER CCD camera (Hamamatsu, Japan). For the life cell imaging, *Arabidopsis* seeds of lines harbouring mutagenized KNL2::KNL2-EYFP/pGWB640 variants or non-mutagenized control were germinated in agar medium in coverslip chambers (Nalge Nunc). Roots growing parallel to the coverslip bottom were analyzed in an LSM 510META confocal microscope (Carl Zeiss) using a 63x oil immersion objective (NA 1.4). EYFP was excited with a 488-nm laser line and fluorescence was recorded with a 505-to 550-nm band-pass filter. Images were analyzed with the LSM software release 3.2.

### Whole-mount preparation

Siliques of different developmental stages were fixed in ethanol-acetic acid (9:1) overnight at 4°C and dehydrated in 70% and 90% ethanol, for 1 h each. The preparation was then cleared in chloral hydrate (chloral hydrate:water:glycerol=8:2:1) overnight at 4°C. Seeds in siliques were counted under a binocular (Carl Zeiss, Germany).

### Cytogenetic techniques

*A. thaliana* inflorescences were fixed in freshly prepared ice-cold ethanol: acetic acid 3:1 and stored at 4°C for chromosome preparations by spreading technique (Armstrong *et al*., 2009). After cell wall digestion, individual buds were dissected on slides, treated with 60% acetic acid, and spread for 30 seconds on a hot plate at 45°C stirring the meiocytes suspension. Post-fixation was done by applying ice-cold 3:1. Air-dried slides were counterstained with DAPI and mounted in Vectashield. Images were acquired in a Nikon Eclipse Ni equipped with a Nikon DS-Qi2 camera and a NIS Elements v. 4.60 software.

### Single Nucleotide Polymorphism (SNP) analysis

Next-Generation Sequencing of genomic DNA was carried out by Eurofins Genomics Europe Shared Services GmbH (Konstanz, Germany) using a Genome Sequencer Illumina NovaSeq 6000 Sequencing System with approximately 5×10^6^ reads for each sample. After adapter trimming with cutadapt (Martin, 2011) version 1.15, next-generation sequence reads were aligned to the TAIR10 assembly with minimap2 (Li, 2018) version 2.17. Alignment records were converted to BAM format with SAMtools (Li *et al*., 2009) and sorted with Novosort (http://www.novocraft.com/products/novosort/). SNP calling was done with BCFtools (Li, 2011) version 1.9 (command ‘mpileup’ and ‘call’) using the parameters ‘-a DP,DV’ to record allelic depths. Only reads with a mapping quality >= Q20 were considered for variant calling. Allelic depths for each sample at bi-allelic SNP sites with a quality score >= Q40 were written to tabular format and read into R (R Core Team, 2017) for further processing. Homozygous genotypes call for the reference (alternative) allele were made if <= 10 % (>= 90 %) of reads supported the variant allele. Heterozygous calls were made if 40-60 % of reads supported the variant allele. If allelic ratios were outside these ranges or the total read depth was < 5, genotype calls were set to missing. SNPs at which the parents carried opposite homozygous alleles were selected to plot graphical genotypes of the progeny along the genome.

### Gene expression analyses

The transcriptome data was retrieved from the *Arabidopsis* RNA-seq Database available on (http://ipf.sustech.edu.cn/pub/athrdb/) (Zhang *et al*., 2020). The gene expression profiles of cenH3, CENP-C, KNL2, and NASP were extracted for different stress treatments including temperature and light stress. Down-regulated treatments among these genes were used for comparative co-expression analysis.

### Flow cytometric ploidy measurements of seeds

To measure the ploidy of seeds, six seeds per pool were chopped together in 500 μl nuclei isolation buffer (Galbraith *et al*., 1983) supplemented with propidium iodide (50 µg/ml) and DNase-free RNase (50µg/ml) in a Petri dish using a sharp razor blade. The resulting nuclei suspensions were filtered a 50 µm mesh (CellTrics, Sysmex-Partec) and measured on a CyFlow Space flow cytometer (Sysmex-Partec), a FACSAria cell sorter (BD Biosciences), or an Influx cell Sorter (BD Biosciences). The presence of haploid/aneuploid seeds within the pools was determined by evaluating the PI fluorescence intensity on a linear scale. Since the precise number of haploids/aneuploids per pool cannot be determined unequivocally, we considered only one seed per pool as being deviating from the diploid status if in addition peak was found. Thus, the number of haploids/aneuploids is an underestimation rather than an overestimation.

