## Supplemental material Fig1-4, Table 2 for "High temperature increases centromere-mediated genome elimination frequency in Arabidopsis deficient in cenH3 or its assembly factor KNL2"

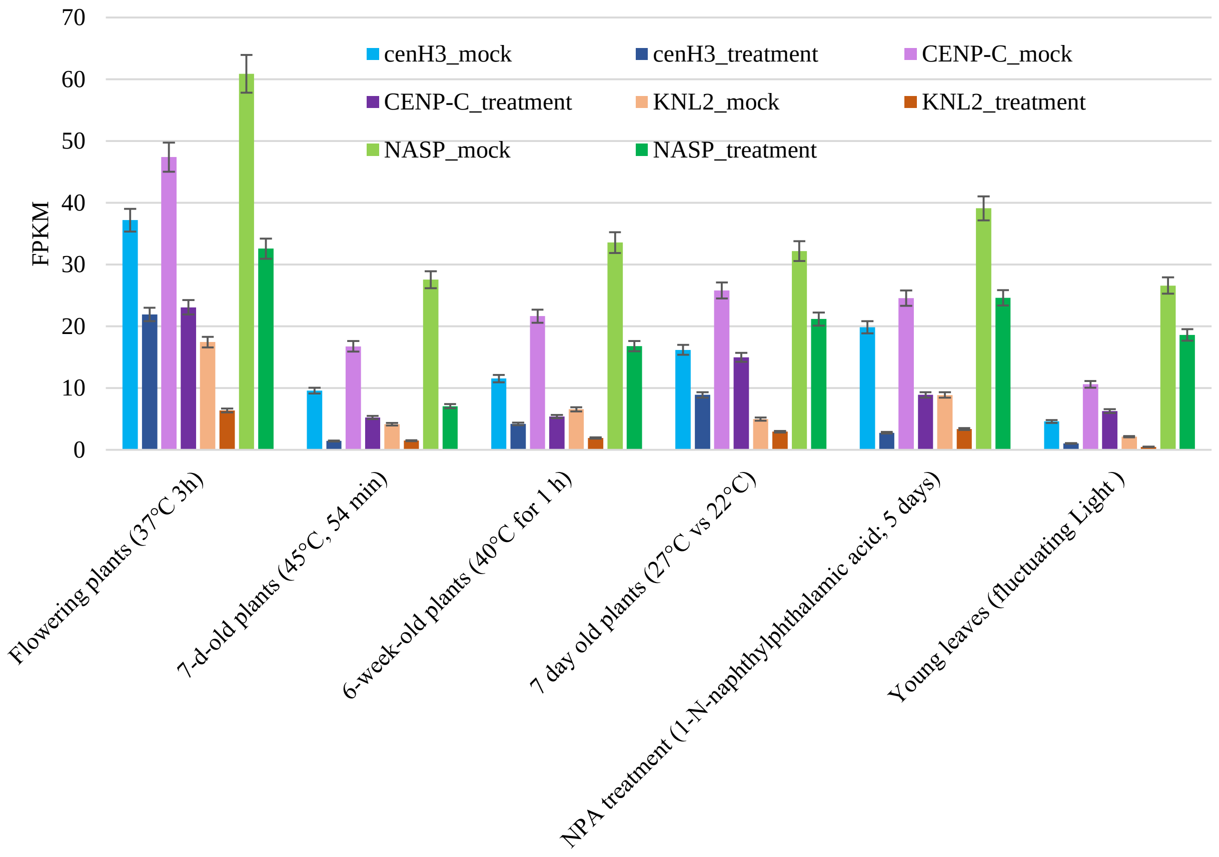


**Figure S1 | The gene expression profiles of cenH3 and kinetochore proteins under different stress treatment in *A. thaliana*.**

Comparative transcriptome analysis showed gene expression difference under different stress treatment in *Arabidopsis*. Heat treatment has leaded downregulated expression of cenH3, CENP-C, KNL2 and NASP. The transcriptional dataset was download under accession numbers: PRJNA363056; PRJDB7363; PRJNA317804; PRJNA497220; PRJNA489360; PRJEB31094.

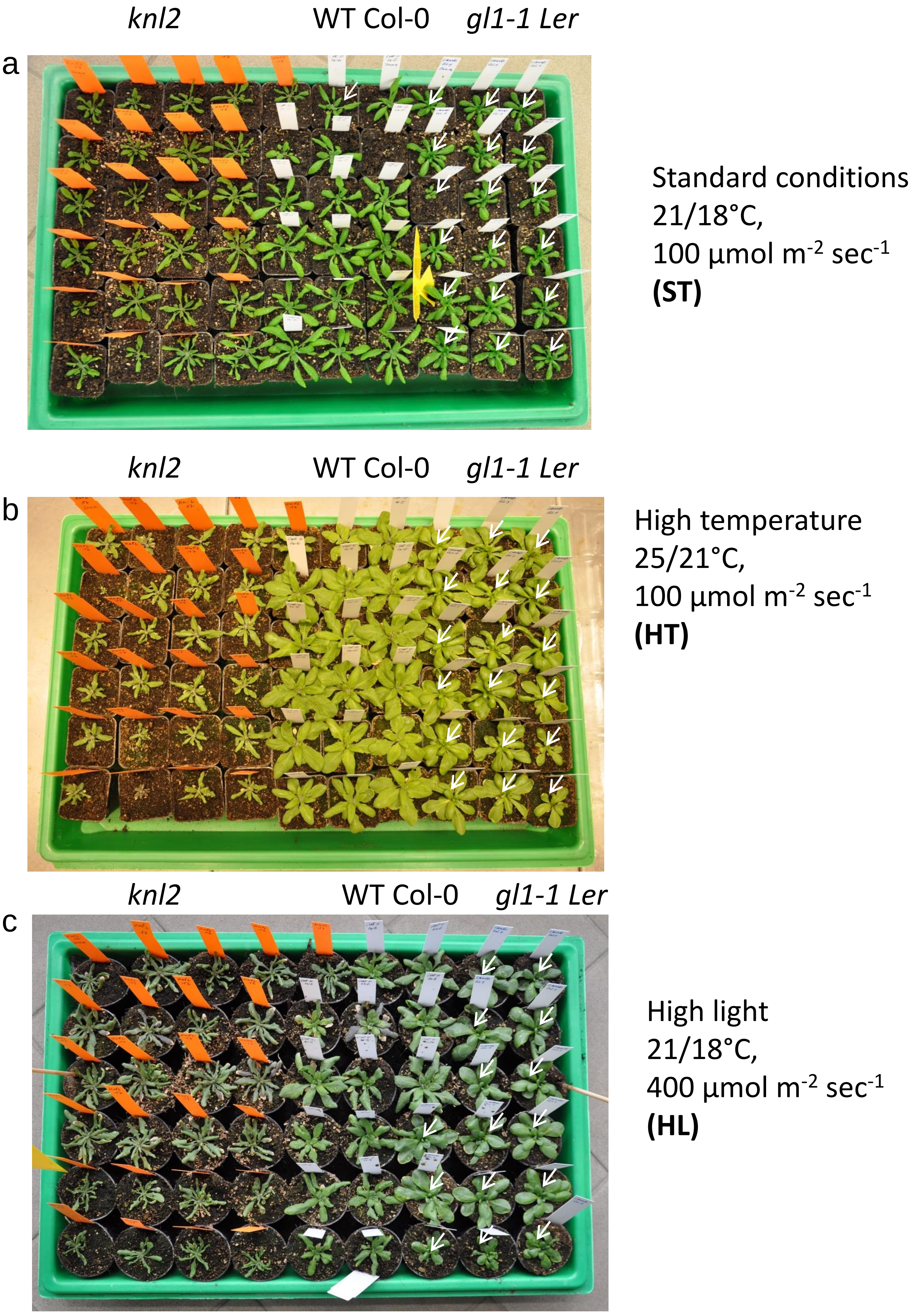

*knl2*
WT Col-0
*gl1-1 Ler*
a
Standard conditions
21/18°C,
100 µmol m$^{-2}$ sec$^{-1}$
**(ST)**
*knl2*
WT Col-0
*gl1-1 Ler*
b
High temperature
25/21°C,
100 µmol m$^{-2}$ sec$^{-1}$
**(HT)**
*knl2*
WT Col-0
*gl1-1 Ler*
c
High light
21/18°C,
400 µmol m$^{-2}$ sec$^{-1}$
**(HL)**

**Figure S2 | Phenotype of 5-week-old *knl2*, wild-type (WT Col-0) and *gl1-1* of *L. erecta* cultivated under different growth conditions.**

All plants were cultivated under standard growth conditions (21-18°C day-night and light intensity 100 µmol $m^{-2}sec^{-1}$) for three weeks, then some plants were left under standard conditions (a) or transferred to higher temperature (25-21°C day night) (b) or higher light intensity (400 µmol $m^{-2}sec^{-1}$) (c) conditions. Orange labels were used to mark *knl2*, while white labels marked *A. thaliana* wild-type Col-0 and the trichome-less *gl1-1* mutant of L. erecta (additionally shown by white arrows).

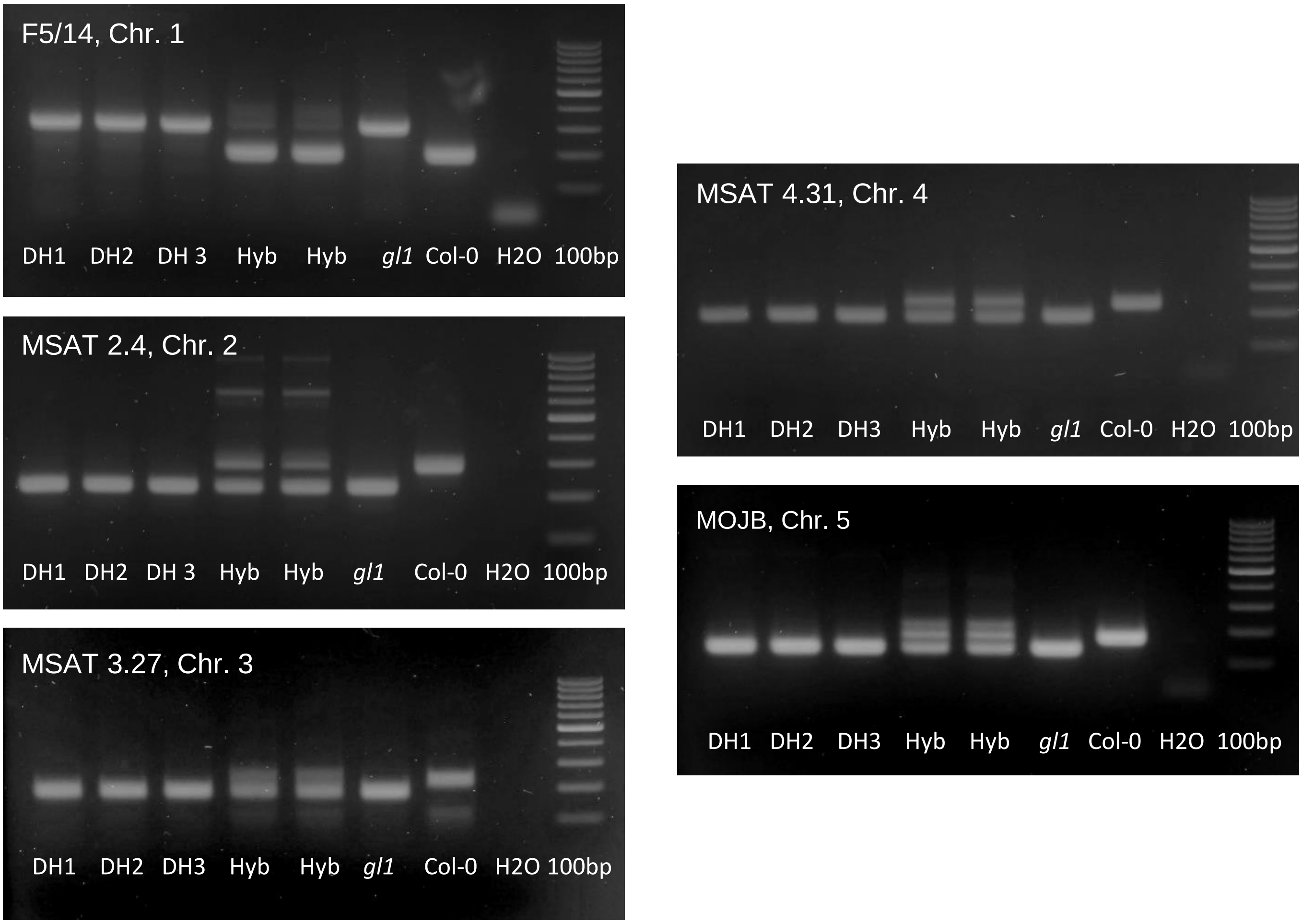


**Figure S3 | Band patterns of chromosome markers amplified on template DNA isolated from double haploids (DH), F1 heterozygous progeny (Hyb) and the parental *gl1-1* (Ler) and Col-0 plants.**

PCR-amplified DNA fragments were electrophoresed on 2.5 % agarose gels, together with size marker Gene Ruler 100bp (Thermo Scientific) on the right side. The chromosome number is shown inside of each panel.

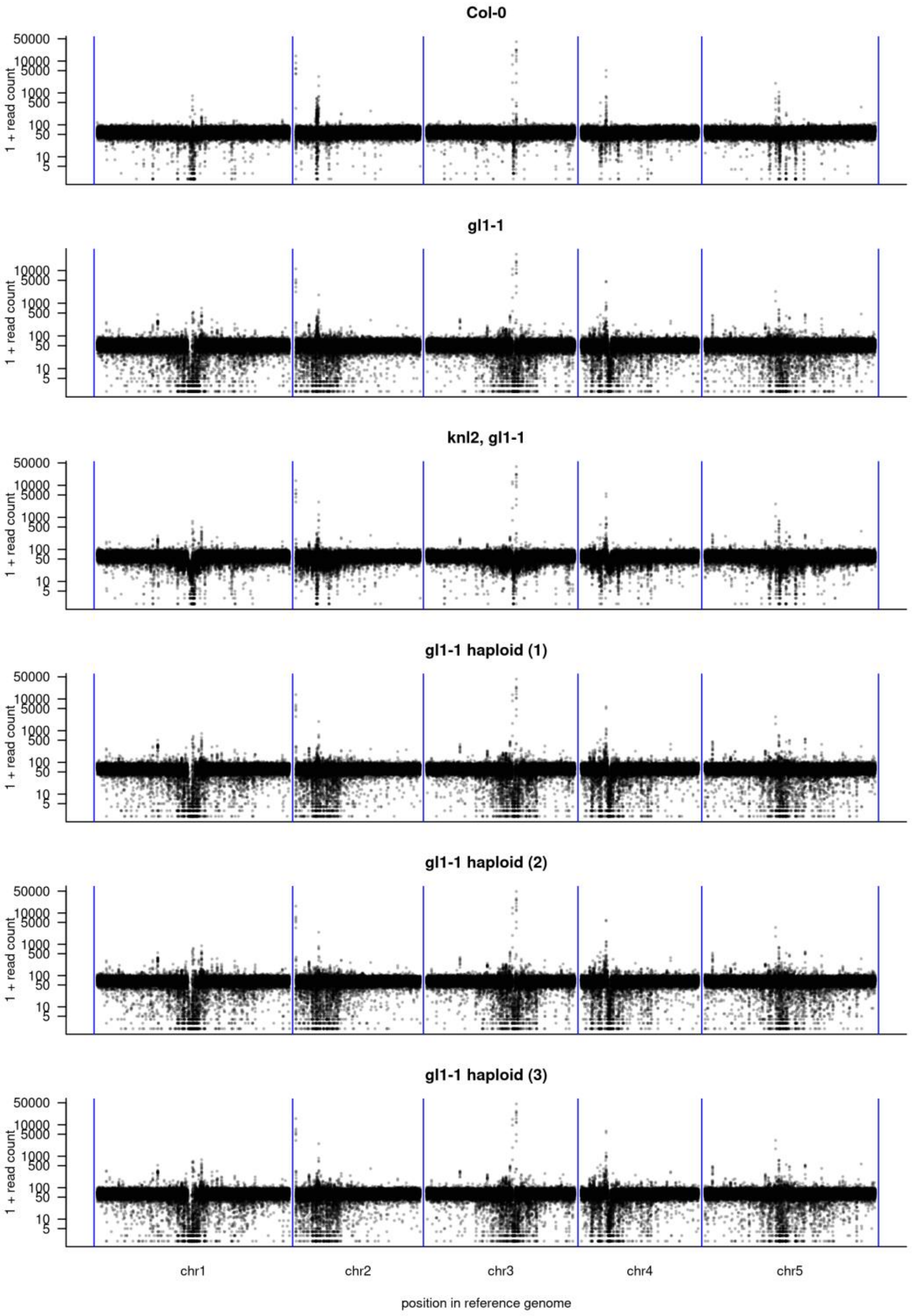


**Figure S4 | Read depth in *gl1-1,* Col-0, a *knl2 / gl1-1* hybrid and three *gl1-1* haploid plants.**

The number of uniquely mapped reads (MAPQ20) in 10 kb windows along the TAIR10 assembly is shown. Due to difficulties in mapping short-reads to repetitive regions coverage around centromeres is variable. No large deletions or duplications private to the *gl1-1* haploids were observed.

**Supplemental Table S2: Primers used in this study**

| Primer name | Primer sequence |
|---|---|
| Gateway cloning | |
| KNL2-attB1gensh | GGGGACAAGTTTGTACAAAAAAGCAGGCTTCAACTATATGATTGTTTACTAC |
| KNL2-attB2 | GGGGACCACTTTGTACAAGAAAGCTGGGTCTTTGATTTTCAAGTTTCTTCG |
| KNL2gen_W_R-f | ATCACTAGAGTTTCGGCGTAACCAAATTCC |
| KNL2gen_W_R-f | GACACAAGCACCCTTCCTGCAAAACATTAA |
| Chromosome markers | |
| F5/14_FP | CTGCCTGAAATTGTCGAAAC |
| F5/14_RP | GGCATCACAGTTCTGATTCC |
| MSAT 2.4_FP | TGGGTTTTTGTGGGTC |
| MSAT 2.4_RP | GTATTATTGTGCTGCCTTTT |
| MSAT 3.27_FP | CCAATGAAAGTTTGATTAC |
| MSAT 3.27_RP | AAATGAAAAATTGAGCTAA |
| MSAT 4.31_FP | AGGGATATGGATTGAGA |
| MSAT 4.31_RP | GCCGTATAACTATTGGTT |
| MOJB_FP | TGAAAGATTTTAGGAGGACAA |
| MOJB_RP | GTAGGAGAAGGGGACAAGTT |
